# Charitability, Compulsion, & the Cost of Control

**DOI:** 10.1101/2025.05.04.652123

**Authors:** Lucila Arroyo, Mimi Liljeholm

## Abstract

Human decision-makers have a well-established preference for free choice. We used a hierarchical gambling task to assess whether this preference extends to environments in which decision outcomes benefit others, specifically charitable organizations. In neurotypical individuals (n=100), we found a reduction in the preference for free choice across self- and charity-benefiting environments when freely chosen options yielded divergent sensory outcome distributions, but not when the outcome distributions of available options were identical – i.e., only when free choice afforded instrumental control. In individuals with obsessive-compulsive traits (n=108) this pattern was reversed, suggesting an unchanged preference for control in charity-benefiting environments. Computational cognitive modeling confirmed a selective reduction in the subjective utility of instrumental control, dissociable from that of free choice, outcome divergence, and monetary reward maximization, in neurotypical individuals gambling for charities. We interpret these results as reflecting cost-benefit analyses weighing the deployment of cognitive effort against the value of control.

## Introduction

Healthy human adults (Bobadilla-Suarez et al., 2017; Bown et al., 2003; Leotti & Delgado, 2011; Leotti et al., 2010; Suzuki, 1997), and a range of non-human animals (Catania, 1975; Perdue et al., 2014; Suzuki, 1999; Voss & Homzie, 1970), have a well-demonstrated preference for choosing freely among options, even when forced selections yield the same reward. This preference may confer a selective advantage: Since the subjective utilities of sensory stimuli (e.g., a particular food or song) fluctuate continuously according to hedonic and homoeostatic dynamics, reward maximization requires an ability to flexibly produce distinct sensory states according to their current utilities (Liljeholm, 2018, 2021, 2022). On the other hand, the computations of second-order statistics required to construct an explicit representation of instrumental control are resource-intense and may not always be adaptive. Here, we assess whether the preference for control extends to prosocial decisions, operationalized as charitable donations, and whether this extension depends on a trait-based need for control.

As an illustration of an environment’s controllability, and its components, consider the scenario depicted in Figure 1A, where each of two gambling ‘rooms’ contain two options, and each option has a probability distribution over three outcome colors, as indicated by the pie charts. Assume that each color can be associated with any amount of monetary loss or gain, and that you must select a room to gamble in *before* those monetary color values are revealed, mimicking the volatility of subjective sensory utilities. In the left room, though afforded free choice, you would have very little control over which color, and thus which monetary outcome, was produced by your selections, since the color distributions are identical across options: Conversely, in the room on the right, though color outcomes diverge across gambling options, such that selection between options can accommodate current color values, you are forced to accept the selections of a computer algorithm, again eliminating your instrumental control. When presented with choice scenarios such as that shown in Figure 1A, knowing that they will get to keep their gambling gains for themselves, neurotypical adults strongly prefer environments that are controllable: i.e., those in which freely chosen actions produce divergent outcome distributions (Liljeholm, 2018, 2021, 2022).

**Fig. 1:**
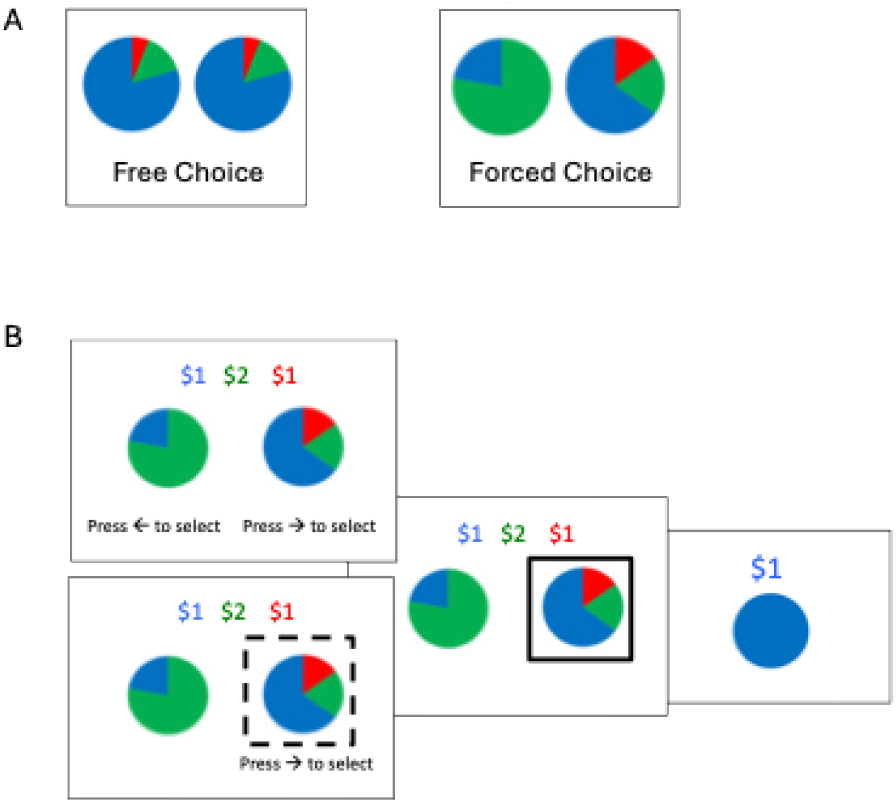
A) Two gambling environments with low (Left) and high (Right) outcome divergence, with neither affording instrumental control. B) Choice, selection & feedback screens on a trial inside a gambling room, illustrating both free (top) and forced (bottom) choice screens. In Forced Choice rooms, dashed lines indicate the required selection.

Note that the assessment of instrumental control requires integration of information both about whether choice is voluntary and whether options yield divergent outcome distributions. Such computations are cognitively demanding, and thus the gain of flexibility must be weighed against the cost of cognitive resources. We hypothesized that prosocial decisions, benefiting charities rather than oneself, might not warrant the deployment of executive functions except, perhaps, in individuals with a compulsive need for control.

### Study 1

Study 1 assessed whether a preference for gambling environments with greater controllability would be reduced when monetary outcomes benefited a charity, self-selected from a list of 60 real charities, rather than oneself. All charitable donations were real.

## Methods

### Participants

The study was a 2 (Free vs. Forced Choice) by 4 (Levels of Outcome Divergence) by 2 (Gamble for oneself or for charity) within-subject design. The sample size was determined by previous work using the same task without the charity conditions (Liljeholm, 2022) to obtain a power of 0.80. The study was administered on Prolific (www.prolific.com), with one hundred participants (32 female, mean age 36.76 ± 11.63) completing the study. Each participant was paid $10 for one hour of participation and could earn up to an additional $36 for themselves, a charity of their choice, or both. All participants gave informed consent and the Institutional Review Board of the University of California, Irvine approved the study.

### Task & Design

Participants were instructed that, in each of several gambling rounds, they would first be required to select a room in which only two slot machines were available, and that they would be restricted to gamble only on those two machines throughout the round, choosing for themselves in free-choice rooms but with the computer alternating between options in forced-choice rooms Participants were further told that the monetary payoffs associated with each slot-machine outcome color would change across rounds and only be revealed once a room had been selected. Finally, participants were instructed that each round would benefit either themselves or charity. The room-choice screen was as shown in Figure 1A, except that individual rooms were labeled as Self-Play or Auto-Play (rather than Forced vs. Free Choice), and that the statement ‘Earnings in this round go to YOU’ or ‘Earnings in this round go to CHARITY’ was printed at the top center of the screen to indicate the nature of the gambling round. Once in a room, the choice, selection, and feedback screens were as illustrated in Figure 1B. In forced-choice rooms (bottom), a dashed square appeared around the computer-selected slot machine, and participants had to press that key in order to proceed through the trial.

The difference between outcome distributions associated with the slot machines inside a room was quantified as the (information theoretic) Jensen-Shannon Divergence between outcome distributions (see methods), with four divergences (0.00, 0.04, 0.15, and 0.20) yielding six unique *differences* in divergence *across* rooms. Each divergence-difference was combined with each possible combination of free and forced choice, for a total of 24 unique room-choice scenarios. Note that only rooms with non-zero divergence *and* free choice are controllable. Unique choice scenarios were repeated across Self-Benefiting and Charity-Benefiting rooms, with different, counterbalanced, color schemes, for 48 total room-selections. Once a room had been selected, four discrete slot-machine choice trials were performed in that room, yielding a total 192 gambling trials. To ensure that stochastically generated monetary payoffs could not account for a preference for greater outcome divergence, reward distributions were constructed such that monetary payoffs were largely balanced and, if anything, biased against hypothesized influences.

Participants were instructed that three gambling rounds would be randomly drawn from all completed rounds and that monetary earnings from the selected rounds would be paid to the participant or their selected charity, according to the nature of the selected rounds. Following the gambling phase, participants were provided with the categories (e.g., children, animals, homelessness) and names of 60 charities and asked to select which should benefit from any charity earnings.

### Computational Cognitive Model

An environment’s controllability may be defined as the IT distance between the outcome probability distributions associated with available actions. Let *P*_*1*_ and *P*_*2*_ be the respective color probability distributions of the two slot machines available in a given room, let *O* be the set of possible color outcomes, and *P(o)* the probability of a particular color outcome. The IT distance is:

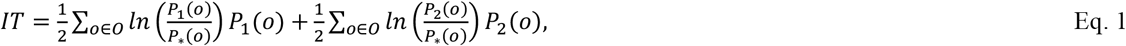

where *ln* is the natural logarithm, yielding nats, and

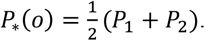

Note that, while an information theoretic divergence can be computed over distributions associated with any number and type of random variable(s), it only reflects instrumental control when computed over sensory outcome distributions associated with freely chosen actions.

An Expected Utility model specified a quantitative integration of the utility of Controllability with conventional monetary reward. First, the acquired value of a particular gambling room (i.e., a particular pair of pie charts) was incrementally updated across gambling rounds, such that

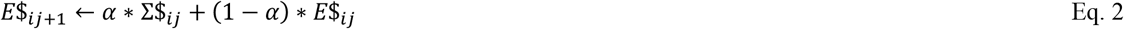

where *E*$_*ij*+1_ is the expected monetary pay-off in room *i* at the start of round *j+1* (i.e., at the start of the next round played in room *i*), Σ$_*ij*_ is the sum of monetary outcomes earned in room *i* at the end of round *j* (i.e., the end of the current round), *E*$_*ij*_ is the expected monetary payoff in room *i* on round *j* (recursively estimated based on the payoff, Σ$_*ij*−1_, and expectation, *E*$_*ij*−1_, in the previous round, *j-1*), and *α* is a free learning rate parameter.

The model further specified three free parameters: *γ*, indicating the subjective utility of Free Choice; *λ*, indicating the subjective utility of the Divergence, *D*, of color outcome distributions associated with the two options (i.e., pie charts) available in a room; and *β* indicating the subjective utility of Controllability (i.e., the combination of Free Choice and non-zero Outcome Divergence)

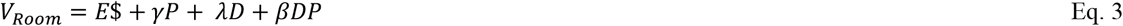

where *P* is an indicator set to 1 for Free Choice and 0 for Forced Choice.

Model-derived room values (*V*_*room*_) were transformed into room choice probabilities using a softmax rule with a noise parameter, *τ*;

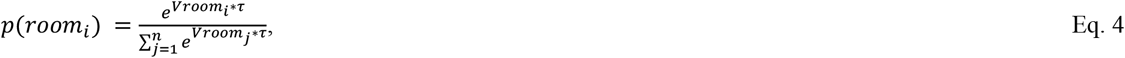

Free model parameters were fit to behavioral data by minimizing the negative log likelihood. All computational variables were implemented using MATLAB (https://www.mathworks.com/).

### Self-Report Measures

Following the charity selection, participants completed a Social Anxiety Questionnaire (SAQ; Lakuta, 2018) and a Test of Self-Conscious Affect (SCA; Tangney, Dearing, Wagner, & Gramzow, 2000), assessing Guilt- and Shame-proneness.

## Results

Recall that only rooms with *both* free choice *and* non-zero outcome divergence afford controllability. In Figure 2A, room-choice preferences are plotted as a function of outcome divergence (0.0 vs. 0.2), free vs. forced choice, and earning for oneself or for a charity. First, when gambling gains were kept for oneself, there was an apparent effect of controllability, such that the preference for rooms with free choice *and* high outcome divergence was greater than the sum of their, also apparent, individual influence, consistent with previous work (Liljeholm, 2022). Second, when gambling gains would instead benefit a charity, the preference for free choice disappeared, while that for greater outcome divergence remained. A 2 (Beneficiary: Self vs. Charity) by 2 (Divergence: Zero vs. Maximal) by 2 (Choice: Free vs. Forced) repeated measures Analysis of Variance performed on the room-choice proportions revealed significant main effects of Choice, F(1,99)=14.53, p<0.001, η^2^_p_ =0.13, and Outcome Divergence, F(1,99)=20.84, p<0.001, η^2^_p_ =0.17, significant Divergence-by-Choice, F(1,99)=13.71, p<0.001, η^2^_p_ =0.12, and Beneficiary-by-Choice, F(1,99)=15.75, p<0.001, η^2^_p_ =0.14, interactions, and a marginally significant Beneficiary-by-Divergence-by-Choice interaction, F(1,99)=3.74, p=0.056, η^2^_p_ =0.04. Despite the significant reduction in the preference for instrumental control when gambling for charities, participants did not differ with respect to monetary reward maximization (earnings relative to that yielded by selection of the best option in each round) across controllable self- and charity-benefiting environments, t(99)=1.46, p=0.148, Cohen’s d=0.146, 95% CI=[-0.052, 0.343].

**Fig. 2:**
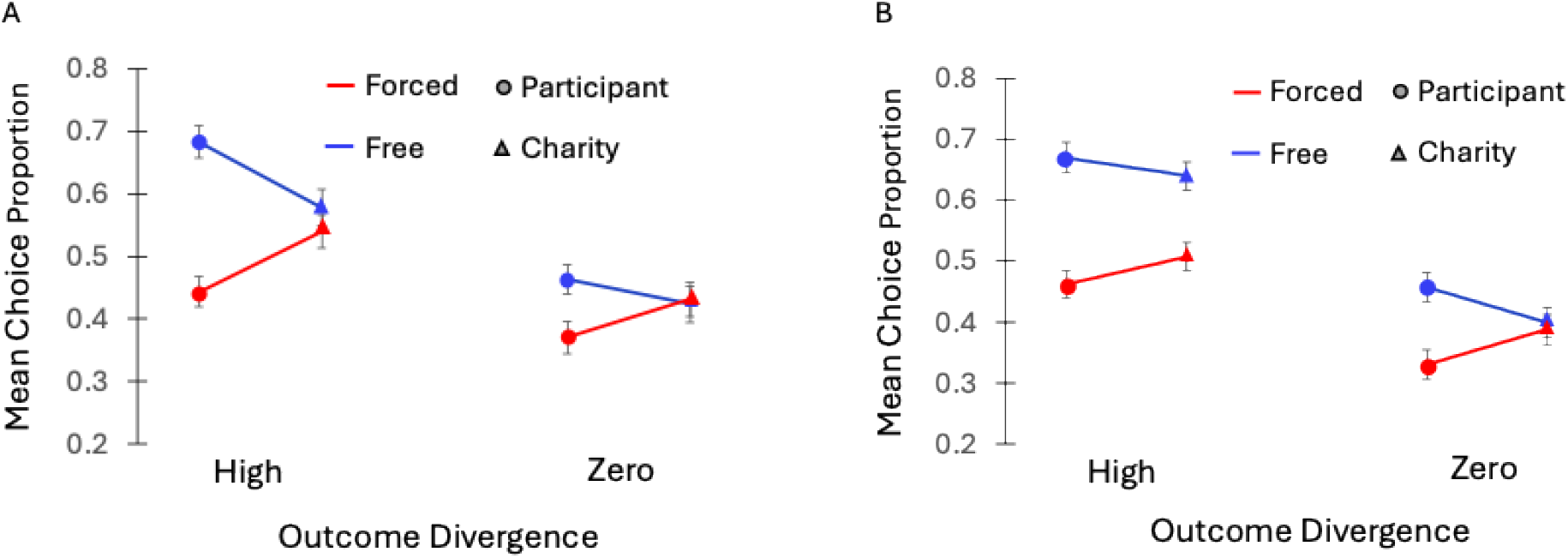
Mean choice proportions for gambling environments, relative to all environments, as a function of High (0.20) vs. Zero Outcome Divergence (the information theoretic distance between outcome distributions across available options), Free vs. Forced choice, and whether the Participant or a Charity benefited from gambling gains. A. Results from Study 1 with neurotypical participants. B. Results from Study 2, with individuals self-identifying as suffering from OCD. Error bars = SEM.

Model-based analyses further confirmed that, beyond a simple reduction in the preference for free-choice, there was a reduction specifically in the preference for instrumental control when gambling for charities. The mean model-derived subjective utilities of Free Choice, Outcome Divergence, and Instrumental Control, across self- and charity earning environments, are plotted in Figure 3 (left side). Planned comparisons revealed significantly reduced utilities of Free Choice, t(99)=2.65, p=0.009, Cohen’s d=0.265, 95% CI=[0.065, 0.464], and Instrumental Control, t(99)=3.45, p<0.001, Cohen’s d=0.345, 95% CI=[0.142, 0.546], but not of Outcome Divergence, t(99)=0.015, p=0.988, Cohen’s d=0.002, 95% CI=[-0.194, 0.198], across self- and charity-benefitting contexts.

**Fig. 3:**
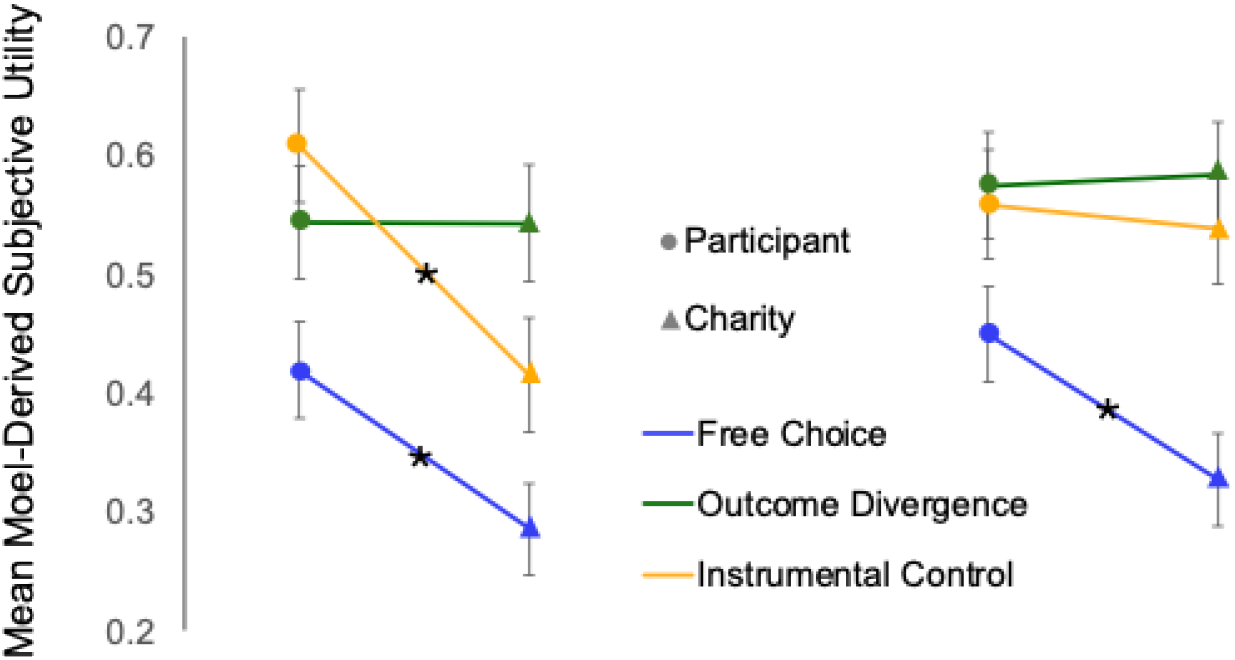
Mean model-derived subjective utilities of Free Choice, Outcome Divergence and Instrumental Control, across participant- and charity-earning conditions, in neurotypical individuals (left) and in individuals self-identifying as suffering from OCD (right). Error bars = SEM. *p<0.01 for two-tailed paired-samples t-tests.

A linear regression with the difference in the model-derived subjective utility of instrumental control across self- and charity-earning environments as the outcome variable found no effects of Social Anxiety, t(99)=-1.02, p=0.313, 95% CI=[-0.018, 0.006], or of proneness towards Shame, t(99)=0.04, p=0.966, 95% CI=[-0.017, 0.018], or Guilt, t(99)=-0.355, p=0.723, 95% CI=[-0.023, 0.016].

### Study 2

Study 1 demonstrated that a well-established preference for environments with greater controllability was significantly reduced when monetary gains benefited a charity rather than oneself. In Study 2, we assessed whether an exaggerated, trait-based, need for control, would counter this reduction. Substantial literature has identified a range of dysregulated agency representations in individuals with obsessive-compulsive disorder (OCD), including both an aberrant sense of agency and an increased desire for control (e.g., Belayachi & Van der Linden, 2010; Borrelli et al., 2024; Moulding & Kyrios, 2007). In Study 2, the study design was identical to that in Study 1, but the task was administered to self-identified individuals with OCD.

## Methods

### Participants

One hundred and eight participants (76 female, mean age 33.19 ± 10.55) were recruited via ResearchMatch, a nonprofit program funded by the National Institutes of Health that helps connect researchers and potential participants – particularly those with medical conditions (https://www.researchmatch.org/). Again, participants were paid $10 for one hour of participation and could earn up to an additional $36 for themselves, a charity, or both.

### Measures

In addition to the SAQ and SCA, participants in Study 2 completed the Dimensional Obsessive-Compulsive Scale (DOCS: Abramowitz et al., 2010) and the Obsessive-Compulsive Inventory Revised (OCI-R: Foa et al., 2002). They were also asked about being diagnosed with OCD by a mental health professional, if they were currently or had ever been in therapy to treat OCD, and if they were currently taking medication to treat OCD. For each question participants could select “Yes”, “No”, or “Rather not say”.

## Results

79.63% of participants had an OCI-R score of 18 or higher, identified as cutoff for diagnosing OCD by Foa et al. (2002), and 79.63% of participants self-reported having been diagnosed with OCD by a mental health professional. Finally, 60.52% of participants had a DOCS score of 21 or higher, identified as a cutoff by Abramowitz et al. (2010). Thus, we concluded the obtained sample suitable for our aim.

Room-choice preferences are plotted in Figure 2B, as a function of outcome divergence (0.0 vs. 0.2), free vs. forced choice, and earning for oneself vs. for a charity. In contrast with the neurotypical results shown in Figure 2A, individuals with OCD do not appear to reduce their preference for free choice in rooms with high outcome divergence when gambling for charities, suggesting an intact utility of instrumental control. An extension of the ANOVA from Study 1 into a 2 (Group: Generic vs. OCD) by 2 (Beneficiary: Self vs. Charity) by 2 (Divergence: Zero vs. Maximal) by 2 (Choice: Free vs. Forced) mixed ANOVA revealed significant main effects of Choice, F(1,206)=37.79, p<0.001, η^2^_p_ =0.16, and Outcome Divergence, F(1,206)=56.99, p<0.001, η^2^_p_ =0.22, and significant Divergence-by-Choice, F(1,206)=28.52, p<0.001, η^2^_p_ =0.12 and Beneficiary-by-Choice interactions, F(1,206)=29.36, p<0.001, η^2^_p_ =0.13, were again obtained, as was a significant Group-by-Beneficiary-by-Divergence-by-Choice interaction, F(1,206)=4.20, p=0.042, η^2^_p_=0.02.

Given the greater number of female participants in the OCD group (76) vs. the Neurotypical group (32), we confirmed that the preference for free choice in rooms with high outcome divergence was greater when earning for oneself than for charity for both female, t(31)=3.04, p=0.005, Cohen’s d=0.538, 95% CI=[0.163,0.906], and non-female t(65)=2.17, p=0.034, Cohen’s d=0.267, 95% CI=[0.020,0.511], Neurotypical subgroups and, conversely, did *not* differ across self- and charity-earning rounds in either female, t(75)=1.59, p=0.117, Cohen’s d=0.182, 95% CI=[-0.045,0.408], or non-female, t(31)=-0.14, p=0.893, Cohen’s d=-0.024, 95% CI=[-0.370,0.323], OCD subgroups. In short, the reduced preference for instrumental control was apparent across these subgroups.

The mean model-derived subjective utilities of Free Choice, Outcome Divergence, and Instrumental Control, across self- and charity earning environments, in individuals with self-reported OCD, are plotted in Figure 3 (right side). Planned comparisons revealed that, contrary to neurotypical participants in Study 1, only the subjective utility of Free Choice, t(107)=2.98, p=0.004, Cohen’s d=0.286, 95% CI=[0.093, 0.478], but *not* that of Instrumental Control, t(107)=0.44, p=0.664, Cohen’s d=0.042, 95% CI=[-0.147, 0.230], nor that of Outcome Divergence, (107)=-0.16, p=0.870, Cohen’s d=-0.016, 95% CI=[-0.204, 0.173], differed significantly across self- and charity-benefitting environments. Thus, OCD individuals differed from their neurotypical counterparts with respect to instrumental control, but not free choice per se.

A linear regression with the difference in the model-derived subjective utility of instrumental control across self- and charity-earning environments as the outcome variable found a significant effect of the OCI-R score, t(107)=- 2.98, p=0.004, 95% CI=[-0.025, -0.005], and a marginally significant effect of Social Anxiety, t(107)=1.79, p=0.076, 95% CI=[-0.001, 0.025], but no effects of the DOCS score, t(107)=1.65, p=0.101, 95% CI=[-0.002, 0.020], or of proneness to Guilt, t(107)=-0.230, p=0.768, 95% CI=[-0.025, 0.019], or Shame, t(107)=-1.40, p=0.164, 95% CI=[-0.027, 0.005]. The selective effect of the OCI-R score confirms that the primary measure used to assess the validity of the sample also predicts variations within the sample.

## Discussion

We used a hierarchical gambling task to assess whether a previously demonstrated preference for instrumental control extends to environments in which decision-outcomes benefit others. Specifically, across gambling environments, we varied access to free choice, the degree to which available options yielded different outcome distributions, and whether gamble earnings benefited the subject or a real-world charity. In Study 1, we found a significant reduction in the preference for free choice across self- and charity-benefiting environments when freely chosen options yielded divergent outcome distributions, but not when the outcome distributions of available options were identical. We interpret these results as reflecting cost-benefit analyses regarding the deployment of cognitive effort (Forbes et al., 2024; Naefgen et al., 2018). Specifically, the computational cost of goal-directed action selection appears to exceed the subjective utility of making charitable donations. At the same time, since free choice is not instrumental in the absence of outcome divergence, it need not tax cognitive resources and thus, in that case, no adjustment in warranted across self- and charity-benefiting conditions.

Intriguingly, in Study 2, when the same task was administered to individuals with obsessive-compulsive disorder (OCD), no reduction in instrumental control across self- and charity-benefiting gambling environments was observed. This is consistent with a rapidly growing literature highlighting a dysregulated sense of agency – the feeling of causing ones’ actions and their sensory consequences (Haggard & Chambon, 2012; Krugwasser et al., 2019) – in individuals with OCD (Borrelli et al., 2024). Several studies have demonstrated lower self-reported agency experiences in individuals with OCD (Gentsch et al., 2012; Giuliani et al., 2017), commonly attributed to a failure to predict and suppress the sensory consequences of their own actions, and a recent meta-analysis of 15 studies concluded that OCD is partially characterized by a dissociation between actual and perceived control (Borrelli et al., 2024).

It should be noted, however, that individuals with OCD differed from neurotypical individuals only with respect to the decision to exercise control across self- and charity-benefiting gambling environments, suggesting that socio-affective dynamics might contribute to the differences between groups. In particular, the seminal ‘inflated responsibility’ model (Salkovskis, 1985), which posits that individuals with OCD have an exaggerated sense of responsibility for negative outcomes, has garnered substantial empirical support (e.g., Mitchell et al., 2020). It is possible that individuals with OCD in the current study felt compelled to ensure that their own lack of effort did not unfavorably influence funds for charitable organizations. More work is needed to better characterize the utility of control in prosocial decision-making, across typical and clinical populations.

## Acknowledgements

This work was supported by a grant from the National Science Foundation (1654187) awarded to M.L.

